## Supplemental figures and tables for "Blind spots on western blots: a meta-research study highlighting opportunities to improve figures and methods reporting"

**Supplementary Figure S1: Unprocessed western blot related to figures 1, 2 and 5.**

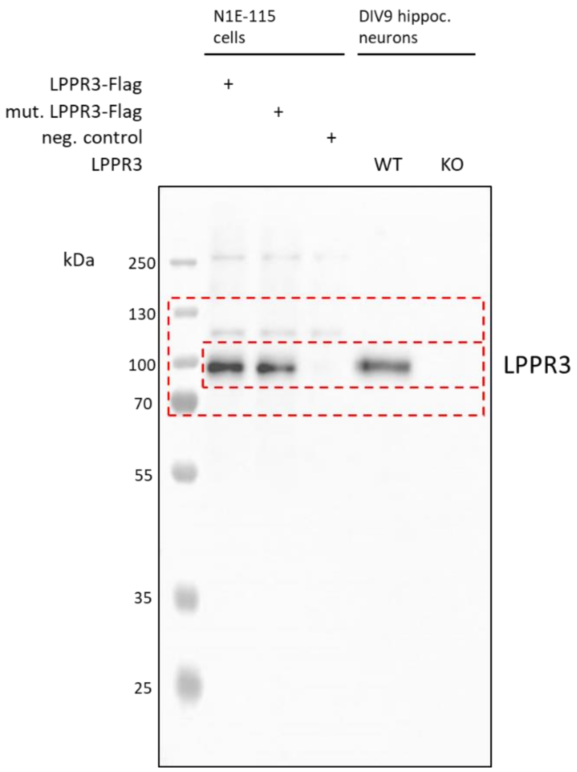


**Method used to capture the image:** The signal was detected with ECL Kit according to manufacturer´s instructions. The blot was incubated in 500 µl ECL reaction for 1 min and imaged using Fusion SL camera (VilberLourmat, Germany) and manufacturer´s software. The blot was imaged in auto-exposure mode with final exposure time of 3 minutes and 25 seconds. Molecular weight marker and chemiluminescent signal images were automatically overlaid by the software creating the image shown here.

For a detailed protocol of materials and methods used to make this western blot image, please see the protocol at dx.doi.org/10.17504/protocols.io.81wgb6z2olpk/v2.

**Supplementary Figure S2: Identification of articles.**

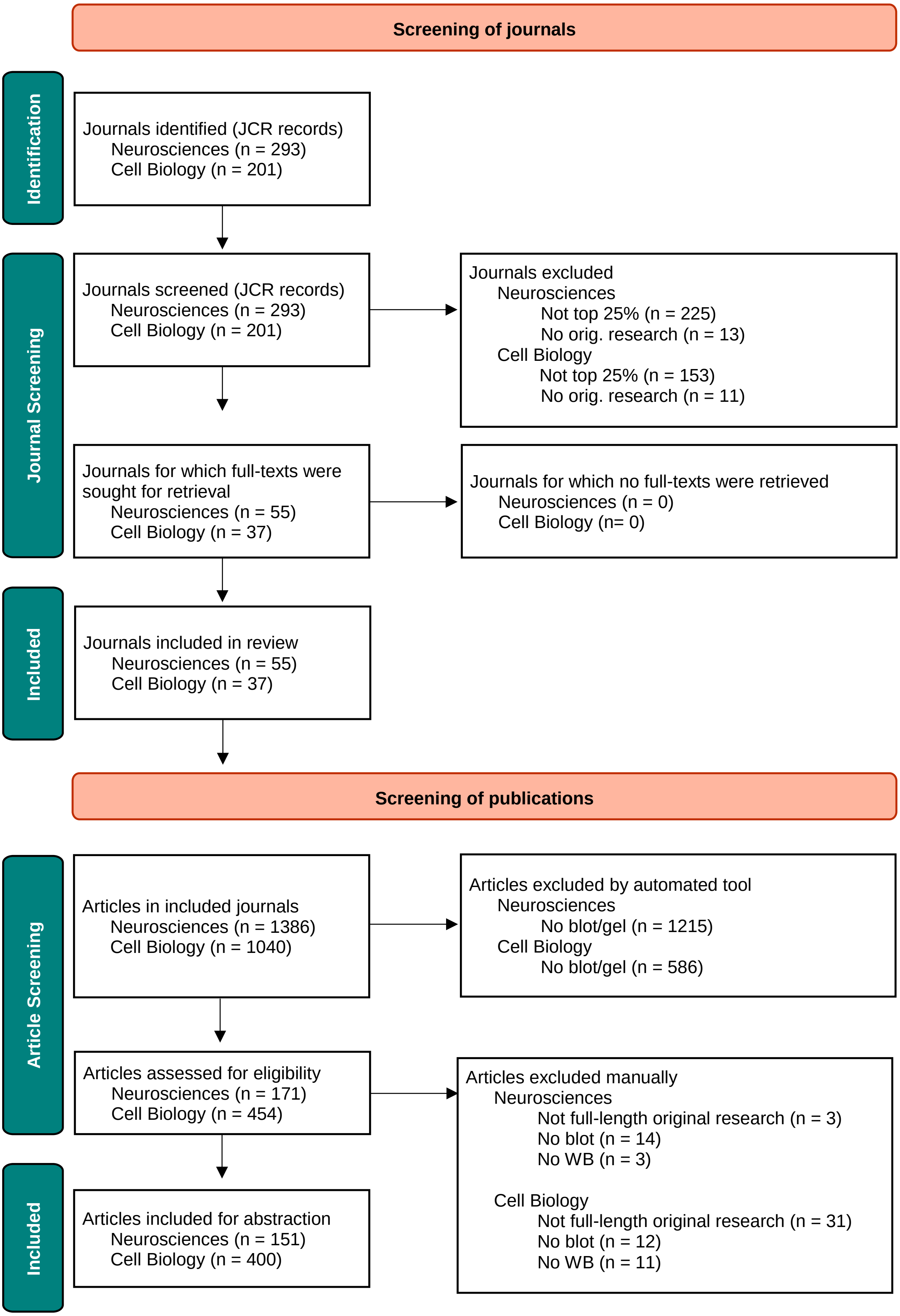

Adapted from:  Page MJ, McKenzie JE, Bossuyt PM, Boutron I, Hoffmann TC, Mulrow CD, et al. The PRISMA 2020 statement: an updated guideline for reporting systematic reviews. BMJ 2021;372:n71. doi: 10.1136/bmj.n71

**Supplementary Table S1**

List of neuroscience journals included in the search, total number of articles screened, the number of articles identified by the tool, and the number and percentage of articles in the study.

| Journal title | Total number of articles screened | Number of articles included by automated tool | Number and percentage of articles included in the study |
|---|---|---|---|
| Acta Neuropathologica | 12 | 6 | 6 (50.00%) |
| Acta Neuropathologica Communications | 9 | 3 | 3 (33.33%) |
| Alzheimer's Research & Therapy | 14 | 1 | 0 (0.00%) |
| Annals of Neurology | 23 | 0 | 0 (0.00%) |
| Behavioral and Brain Sciences | 1 | 0 | 0 (0.00%) |
| Biological Psychiatry | 32 | 2 | 2 (6.25%) |
| Biological Psychiatry Cognitive Neuroscience and Neuroimaging | 16 | 0 | 0 (0.00%) |
| Bipolar Disorders | 11 | 0 | 0 (0.00%) |
| Brain | 38 | 2 | 2 (5.26%) |
| Brain Behavior and Immunity | 31 | 4 | 4 (12.90%) |
| Brain Pathology | 23 | 2 | 2 (8.70%) |
| Brain Stimulation | 19 | 0 | 0 (0.00%) |
| Cephalalgia | 11 | 0 | 0 (0.00%) |
| Cerebral Cortex | 59 | 7 | 5 (8.47%) |
| Cerebrovascular and Brain Metabolism Reviews | 15 | 2 | 2 (13.33%) |
| Cognitive Computation | 18 | 0 | 0 (0.00%) |
| Developmental Cognitive Neuroscience | 11 | 0 | 0 (0.00%) |
| European Journal of Neurology | 70 | 0 | 0 (0.00%) |
| Fluids and Barriers of the CNS | 6 | 0 | 0 (0.00%) |
| Frontiers in Aging Neuroscience | 102 | 9 | 8 (7.84%) |
| Frontiers in Cellular Neuroscience | 47 | 6 | 6 (12.77%) |
| Frontiers in Molecular Neuroscience | 28 | 8 | 6 (21.43%) |
| Glia | 13 | 7 | 6 (46.15%) |
| Journal of Neural Engineering | 28 | 0 | 0 (0.00%) |
| Journal of Neurochemistry | 25 | 8 | 6 (24.00%) |
| Journal of Neuroinflammation | 23 | 15 | 14 (60.87%) |
| Journal of Neuroscience | 83 | 11 | 10 (12.05%) |
| Journal of Pain | 10 | 0 | 0 (0.00%) |
| Journal of Parkinson's Disease | 35 | 3 | 2 (5.71%) |
| Journal of Pineal Research | 3 | 1 | 1 (33.33%) |
| Journal of Psychiatry and Neuroscience | 10 | 0 | 0 (0.00%) |
| Molecular Autism | 5 | 1 | 0 (0.00%) |

| | | | |
|---|---|---|---|
| Molecular Neurobiology | 37 | 21 | 21 (56.76%) |
| Molecular Neurodegeneration | 7 | 2 | 2 (28.57%) |
| Molecular Psychiatry | 38 | 7 | 7 (18.42%) |
| Multiple Sclerosis Journal | 20 | 0 | 0 (0.00%) |
| Nature Human Behaviour | 22 | 0 | 0 (0.00%) |
| Nature Neuroscience | 11 | 2 | 2 (18.18%) |
| Neural Networks | 34 | 2 | 0 (0.00%) |
| NeuroImage | 77 | 3 | 0 (0.00%) |
| Neurobiology of Disease | 14 | 4 | 4 (28.57%) |
| Neurobiology of Stress | 12 | 3 | 3 (25.00%) |
| Neurology Neuroimmunology & Neuroinflammation | 16 | 0 | 0 (0.00%) |
| Neuron | 29 | 1 | 1 (3.45%) |
| Neuropathology and Applied Neurobiology | 12 | 6 | 6 (50.00%) |
| Neuropsychopharmacology | 37 | 1 | 1 (2.70%) |
| Neuroscience & Biobehavioral Reviews | 36 | 0 | 0 (0.00%) |
| Neurotherapeutics | 20 | 8 | 8 (40.00%) |
| Pain | 38 | 2 | 2 (5.26%) |
| Progress in Neurobiology | 14 | 3 | 3 (21.43%) |
| Sleep | 33 | 0 | 0 (0.00%) |
| The Journal of Headache and Pain | 25 | 3 | 2 (8.00%) |
| Translational Neurodegeneration | 3 | 1 | 1 (33.33%) |
| Translational Stroke Research | 8 | 2 | 2 (25.00%) |
| npj Parkinson's Disease | 12 | 2 | 1 (8.33%) |

**Supplementary Table S2**

List of cell biology journals included in the search, total number of articles screened, the number of articles identified by the tool, and the number and percentage of articles in the study.

| Journal title | Total number of articles screened | Number of articles included by automated tool | Number and percentage of articles included in the study |
|---|---|---|---|
| Aging Cell | 17 | 7 | 7 (41.18%) |
| American Journal of Respiratory Cell and Molecular Biology | 16 | 7 | 7 (43.75%) |
| Autophagy | 20 | 8 | 8 (40.00%) |
| Cancer Cell | 18 | 3 | 3 (16.67%) |
| Cell | 36 | 11 | 8 (22.22%) |
| Cell Calcium | 8 | 0 | 0 (0.00%) |
| Cell Death & Differentiation | 15 | 11 | 10 (66.67%) |
| Cell Death & Disease | 98 | 82 | 81 (82.65%) |
| Cell Discovery | 11 | 3 | 1 (9.09%) |
| Cell Metabolism | 16 | 3 | 3 (18.75%) |
| Cell Proliferation | 16 | 10 | 10 (62.50%) |
| Cell Reports | 134 | 66 | 56 (41.79%) |
| Cell Research | 13 | 4 | 3 (23.08%) |
| Cell Stem Cell | 8 | 1 | 1 (12.50%) |
| Cell Systems | 3 | 0 | 0 (0.00%) |
| Cellular and Molecular Life Sciences | 25 | 4 | 3 (12.00%) |
| Current Biology | 90 | 10 | 8 (8.89%) |
| Developmental Cell | 17 | 6 | 6 (35.29%) |
| EMBO Reports | 35 | 15 | 14 (40.00%) |
| Genes & Development | 11 | 6 | 6 (54.55%) |
| Journal of Biomedical Science | 5 | 4 | 4 (80.00%) |
| Journal of Cell Biology | 29 | 18 | 12 (41.38%) |
| Journal of Extracellular Vesicles | 12 | 4 | 4 (33.33%) |
| Matrix Biology | 2 | 1 | 1 (50.00%) |
| Molecular Cell | 31 | 19 | 13 (41.94%) |
| Nature Cell Biology | 18 | 6 | 5 (27.78%) |
| Nature Medicine | 38 | 3 | 0 (0.00%) |
| Nature Structural & Molecular Biology | 20 | 4 | 2 (10.00%) |
| Oncogene | 55 | 43 | 41 (74.55%) |
| Protein & Cell | 6 | 3 | 2 (33.33%) |
| Science Signaling | 12 | 7 | 6 (50.00%) |
| Science Translational Medicine | 27 | 17 | 17 (62.96%) |
| Signal Transduction and Targeted Therapy | 45 | 13 | 5 (11.11%) |
| Stem Cell Reports | 23 | 6 | 5 (21.74%) |

| | | | |
|---|---|---|---|
| Stem Cell Research & Therapy | 59 | 30 | 29 (49.15%) |
| The EMBO Journal | 23 | 13 | 13 (56.52%) |
| The Plant Cell | 28 | 6 | 6 (21.43%) |

**Supplementary Table S3. Antibody reporting template.**

| | Type | Name | Source | Catalog nr | Lot nr | RRID | Dilution | Application | Which secondary was used (nr)? |
|---|---|---|---|---|---|---|---|---|---|
| 1 | Primary | Turbo-GFP | Evrogen | AB513 | 51301010912 | AB_20544089 | 1:2000 | WB | 3 |
| | | | | | | | 1:100 | IHC | 5 |
| 2 | Primary | NT5D1 | Thermo Fisher Scientific | MA5-25214 | TH0923445 | AB_2723416 | 1:1000 | WB | 4 |
| 3 | Secondary | anti-rabbit-HRP | Innovative Research | IGAR-HRP | 150943 | AB_11041560 | 1:5000 | WB | NA |
| 4 | Secondary | anti-mouse-HRP | Fitzgerald Industries International | 43R-IG067hrp | 394573872 | AB_1287465 | 1:10 000 | WB | NA |
| 5 | Secondary | anti-mouse-Alexa488 | ThermoFisher Scientific | A97457 | W836911 | AB_2333901 | 1:500 | IHC | NA |
| 6 | | | | | | | | | |
| 7 | | | | | | | | | |
| 8 | | | | | | | | | |
